# Probing RNA conformational equilibria within the functional cellular context

**DOI:** 10.1101/634576

**Authors:** Laura R. Ganser, Chia-Chieh Chu, Hal P. Bogerd, Megan L. Kelly, Bryan R. Cullen, Hashim M. Al-Hashimi

**Affiliations:** Department of Biochemistry; Duke University Medical Center; Durham, NC, 27710; USA; Department of Molecular Genetics and Microbiology, Center for Virology; Duke University Medical Center; Durham, NC, 27710; USA

**Keywords:** HIV-1 transactivation response element, TAR, Rev response element, RRE, RNA dynamics, RNA drug discovery, transactivation, excited state, RNA structure, structure mapping, cellular activity, conformational switches

## Abstract

Many regulatory RNAs undergo changes in their structure from the dominant ground-state (GS) toward short-lived low-abundance ‘excited-states’ (ES) that reorganize local elements of secondary structure. ESs are increasingly observed *in vitro* and implicated in the folding and biological activities of regulatory RNAs and as targets for developing therapeutics. However, whether these ESs also form with comparable abundance within the complex cellular environment remains unknown. Here, we developed an approach for assessing the relative stability and abundance of RNA ESs within the functional cellular context. The approach uses point substitution mutations to increase the population of an inactive ES relative to the active GS. The cellular activity of such ES-stabilizing mutants then provides an indirect measure of any residual population of the active GS within the functional cellular context. Compensatory rescue mutations that restore the GS are used to control for changes in cellular activity arising due to changes in sequence. The approach is applied to probe ESs in two highly conserved and functionally important regulatory RNAs from HIV-1: the transactivation response element (TAR) and the Rev response element (RRE). For both RNAs, ES-stabilizing mutations inhibited cellular activity to a degree that correlates with the extent to which they stabilize the ES relative to the GS *in vitro*. These results indicate that the non-native ESs of TAR and RRE likely form in cells with abundances comparable to those measured *in vitro* and their targeted stabilization provides a new avenue for developing anti-HIV therapeutics.

## Main Text

RNA is a highly flexible biomolecule and understanding its folding behavior and biological activity requires that it is described not as a static structure, but rather as a dynamic ensemble of many interconverting conformations with variable populations and lifetimes (Leulliot and Varani, 2001; Cruz and Westhof, 2009; Mustoe et al., 2014; Schroeder, 2018; Ganser et al., 2019). The relative populations of different conformations in the ensemble change during RNA folding or as the RNA executes its biological activity(Kim and Breaker, 2008; Cruz and Westhof, 2009; Dethoff et al., 2012a; Abeysirigunawardena et al., 2017; Ganser et al., 2019). Such changes in the RNA ensemble help optimize interactions with protein and ligand binding partners(Williamson, 2000), provide a basis for altering RNA activity in RNA-based molecular switches(Breaker, 2011; Fürtig et al., 2015), and satisfy the distinct structural requirements of the multi-step catalytic cycles of ribozymes and ribonucleoprotein machines(Zhuang et al., 2000, 2002). Furthermore, disrupting the RNA ensemble is a proven strategy for targeting RNA therapeutically(Walter et al., 1999; Hermann, 2002; Stelzer et al., 2011; Dibrov et al., 2014; Ganser et al., 2018). Therefore, methods to characterize dynamic ensembles of RNA are critical for understanding RNA folding behavior, cellular activity, and for extending rational structure-based drug discovery approaches to target highly flexible RNAs.

A ubiquitous feature of RNA ensembles is short-lived and low-abundance conformations often referred to as ‘excited-states’ (ESs)(Skrynnikov et al., 2002; Xue et al., 2015). Relative to the dominant ground-state (GS), ESs feature localized changes in secondary structure in and around non-canonical motifs such as bulges and internal loops(Dethoff et al., 2012b; Xue et al., 2015). RNA ESs have been observed for a variety of regulatory RNAs and are increasingly implicated in the folding(Xue et al., 2016) and biological activities of RNAs(Helmling et al., n.d.; Kimsey et al., 2015; Zhao et al., 2017). Because ESs often disrupt structural elements required for function, they are also potentially unique and attractive drug targets(Dethoff et al., 2012b; Chu et al., 2019). However, it remains to be determined whether RNA ESs form in cells with similar abundance as observed *in vitro*. This knowledge is important for verifying the roles for RNA ESs within cells and to establish their validity as viable RNA drug targets(Dethoff et al., 2012b; Connelly et al., 2016).

There have been substantial advances in methods to characterize various aspects of the RNA dynamic ensemble under *in vitro* solution conditions that combine a variety of experimental measurements with computational modeling(Salmon et al., 2014; Xue et al., 2015; Shi et al., 2016; Tian and Das, 2016; Nichols et al., 2018). However, probing RNA dynamics *in vivo* remains an unmet challenge(Rogers and Heitsch, 2014; Li and Aviran, 2017; Woods et al., 2017; Spasic et al., 2018). Recent studies using transcriptome-wide structure probing experiments indicate that the cellular environment can affect RNA folding relative to *in vitro*(Beaudoin et al., 2018; Mustoe et al., 2018; Sun et al., 2019). For example, while G-rich sequences from mammalian RNA stably form thousands of G-quadruplexes *in vitro*, they were shown to be overwhelmingly unfolded *in vivo*(Guo and Bartel, 2016). Such results underscore the importance of measuring dynamic ensembles of RNAs within cells and ideally at the specific time and place where a given RNA executes its function. Despite some advances in using reactivity data to interpret RNA ensembles(Rogers and Heitsch, 2014; Li and Aviran, 2017; Woods et al., 2017; Spasic et al., 2018), the dependence of reactivity on structure is not entirely understood(Rogers and Heitsch, 2014; Spasic et al., 2018). This combined with low sensitivity to low-abundance transient structures such as ESs has made it difficult to probe secondary structural conformational equilibria within cells(Ganser et al., 2019). In addition, such approaches do not probe RNA structure at a specific time and place where a given RNA function is executed within the cell, but rather, the reactivity data can reflect the structure of the RNA in many in different contexts.

We developed an approach to assess the abundance and energetic stability of low-abundance short-lived ESs relative to the GS in cells (Fig 1). The approach uses point substitution mutations, which are expected to minimally impact the biological activity of the GS, to invert the conformational equilibrium so that the inactive ES is the dominant state *in vitro* which can be determined using NMR spectroscopy. The cellular activity (e.g. transcriptional activation) of such ES-stabilizing mutants then provides an indirect measure of any residual population of the active GS within the functional cellular context (Fig 1). If an ES forms in cells with abundance comparable to that measured *in vitro*, mutants that stabilize the ES should inhibit downstream cellular activity to a degree commensurate with the degree to which they decrease the GS population *in vitro* (Fig 1). Alternatively, if the cellular environment preferentially biases the ensemble away from the ES in favor of the GS relative to the *in vitro* environment, the residual population of the GS will be greater than expected based on *in vitro* measurements, and the impact of the mutation on activity will be attenuated in cells (Fig 1). Key to the approach is the use of compensatory rescue mutations that restore the GS conformation while maintaining the original mutation to control for any changes in cellular activity arising due to changes in the RNA sequence. In this manner, we can probe how the cellular environment affects RNA secondary structural equilibra involving exceptionally low-abundance (populations <1%) conformational states within the functional cellular context. The approach assumes that the contribution of the inactive ES to the measured functional readout is negligible and that the observed cellular activity is determined by the abundance of the GS relative to the inactive ES and not their kinetics of inter-conversion.

**Figure 1.**
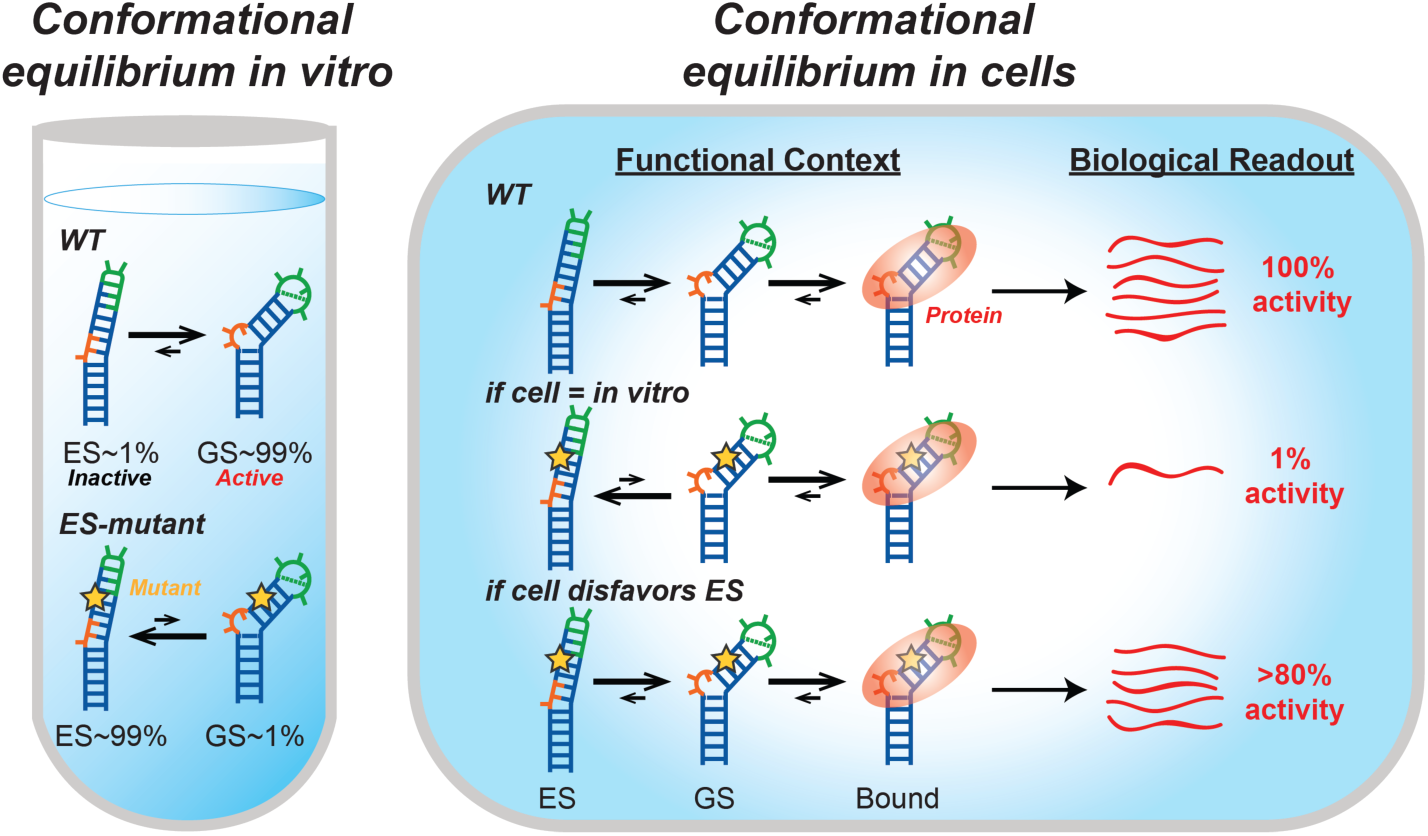
Evaluating the impact of the cellular environment on conformational equilibria using targeted stabilization of RNA ESs and functional readouts to measure the population of the active RNA GS. Substitution mutations (shown as star) redistribute the WT RNA ensemble *in vitro*, increasing the abundance of an ES over the GS to various degrees measured by NMR spectroscopy. The impact of the mutations on cellular activity is then measured using an appropriate assay.

We applied the above approach to the trans-activation response element (TAR), a highly structured and conserved regulatory RNA element in HIV-1 (Fig 2). TAR is located at the 5’ end of the HIV-1 viral genome where it promotes transcription elongation by binding to the viral protein Tat and human super elongation complex (SEC)(Wei et al., 1998; Bieniasz et al., 1999; Ivanov et al., 2000; Kim et al., 2002; Fujinaga et al., 2004; He et al., 2010; Sobhian et al., 2010). This interaction is required to release RNA polymerase II from promoter-proximal pausing, allowing for efficient transcription of the viral genome(He et al., 2010). The TAR GS structure is critical for its biological activity, and Tat and SEC form extensive interactions with the TAR bulge, upper stem, and apical loop(Puglisi et al., 1995; Schulze-Gahmen et al., 2016; Pham et al., 2018; Schulze-gahmen and Hurley, 2018; Chavali et al., 2019). TAR has previously been shown to adopt two ESs, termed ‘ES1’(Dethoff et al., 2012b) and ‘ES2’(Lee et al., 2014) (Fig 2A). ES1 forms through conformational changes that zip up the apical loop creating two non-canonical base pairs (bps) (Fig 2A)(Dethoff et al., 2012b). On the other hand, ES2 remodels the entire secondary structure from the bulge motif to the apical loop, creating a 1-nucleotide bulge, a dinucleotide apical loop, and four non-canonical mismatches (Fig 2A)(Lee et al., 2014). Because ES2 greatly remodels the TAR structure, it is highly unlikely to support transcriptional trans-activation. Therefore, ES2-stabilizing mutations are expected to significantly inhibit viral transcription unless the cellular environment substantially biases the equilibrium to favor the GS.

**Figure 2.**
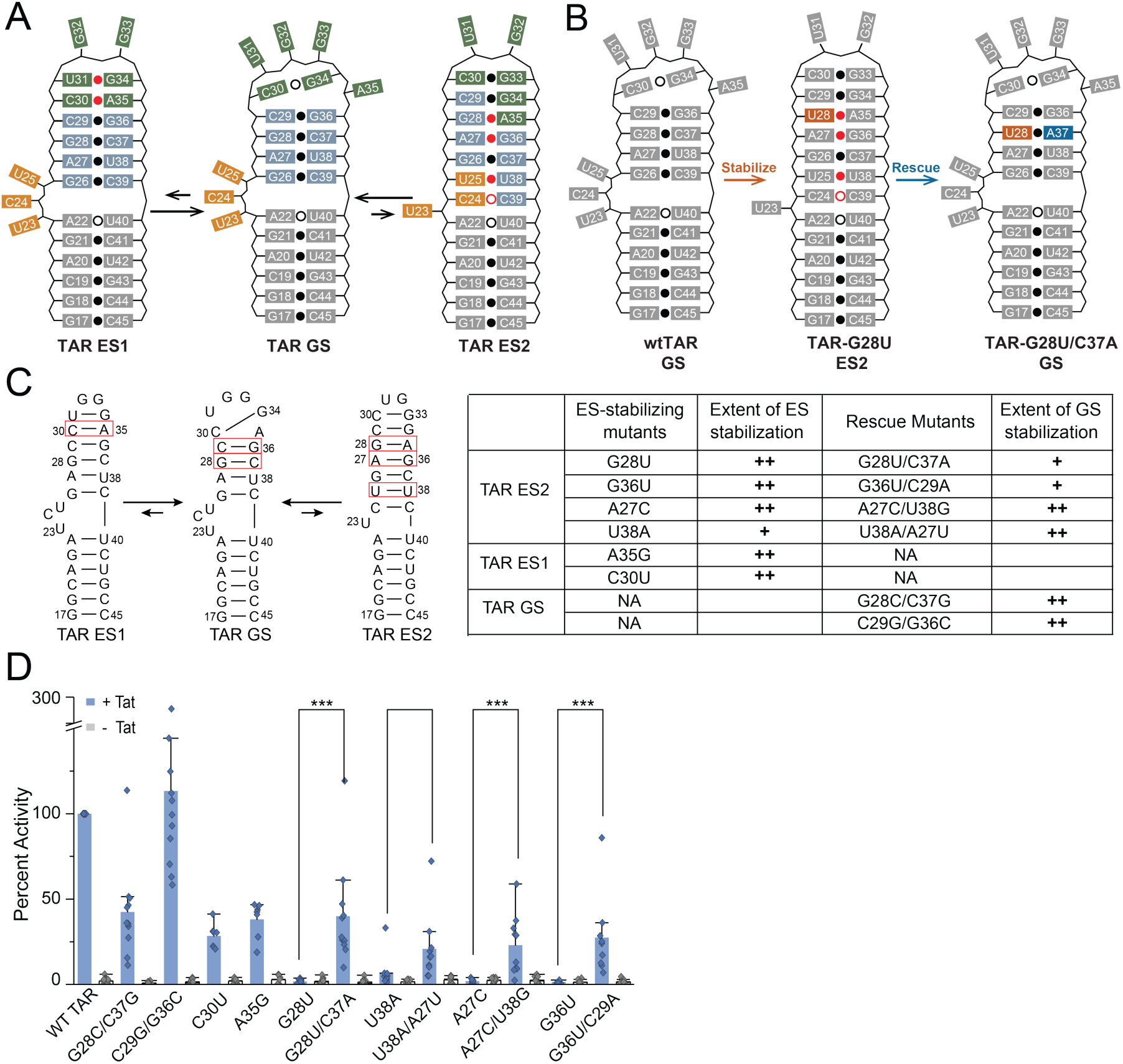
Probing the conformational equilibrium of HIV-1 TAR *in vitro* and *in vivo*. **A.** HIV-1 TAR RNA has two excited states that alter base pairing in the bulge (orange), upper stem (blue), and apical loop (green). Weak base pairs are indicated with an open dot and non-Watson-Crick base pairs are indicated with a red dot. **B.** Example of the stabilize-and-rescue approach for trapping TAR ES2 using the ES-stabilizing mutant G28U (orange) and its corresponding rescue mutation C37A (blue). Weak base pairs are indicated with an open dot and non-Watson-Crick base pairs are indicated with a red dot. **C.** Summary of all TAR mutants. Base pairs that were mutated are highlighted (red box). The extent of stabilization was estimated based on NMR line broadening (Supplementary Figs 2-4). (++) minimal line broadening, (+) partial line broadening, and (-) extensive line broadening. **D.** Tat-dependent trans-activation assay for TAR mutants. HeLa cells were transfected with pFLuc-TAR reporter plasmid and an RLuc internal control in the presence (+ Tat) or absence (-Tat) of a Tat expression plasmid. At 48 hr post transfection, cell lysates were tested for luciferase activity. Reported values are the quotient of FLuc and RLuc activities with values normalized to WT TAR for every independent replicate reported as the mean ± SD. Statistical significance was determined after log-transformation of the data; *P < 0.05, **P < 0.01, ***P < 0.001 (two-sided ANOVA).

We designed a series of mutations that invert the conformational equilibrium so that ES2 is the dominant conformation and corresponding compensatory secondary mutations that restore the GS conformation to control for changes in cellular activity arising purely due to changes in the RNA sequence (Fig 2B and 2C, Supplementary note 1, and Supplementary Figs 1-3). It has previously been shown(Lee et al., 2014) that the TAR-G28U mutant forms ES2 as the dominant conformation. The mutation replaces a mismatch in the ES with a more stable Watson-Crick bp, while simultaneously replacing a Watson-Crick bp in the GS with a mismatch (Fig 2B). We designed the mutant TAR-G28U/C37A, which has a compensatory rescue mutation so that the net effect is replacing the Watson-Crick G28-C37 bp in the GS, with another Watson-Crick U28-A37 bp (Fig 2B). To test the robustness of the results and to control for minor effects of the mutation on the GS, we used this approach to design a total of four ES2-stabilizing mutants and corresponding rescue mutations. Additionally, we generated two positive controls, TAR-G36C/C29G and TAR-G28C/C37G, which invert a Watson-Crick bp in the GS and are therefore expected to maintain the GS conformation.

Key to our approach is the use of NMR spectroscopy (see Methods, Fig 2C, Supplementary Fig 2) to assess the degree to which a given mutation biases the ensemble toward the GS or ES conformation *in vitro* as this sets the baseline for the expected biological activity within cells. If a mutation stabilizes the ES to 90%, we would expect the mutant to have 10% residual activity relative to wild-type, whereas if the mutation stabilizes the ES to 99%, we would expect 1% activity and so on. Based on NMR relaxation dispersion experiments(“Characterizing micro-to-millisecond chemical exchange in nucleic acids using off-resonance R1ρ relaxation dispersion,” n.d.; Sekhar and Kay, 2013; Xue et al., 2015), TAR-G28U stabilizes ES2 to a population of ∼99.8% (Fig 2C, Supplementary Fig 4, and Supplementary Tables 1 and 2). The experiments show that TAR-G28U back-exchanges with a GS-like conformation that has an equilibrium population of ∼0.2% (Supplementary Fig 4 and Supplementary Tables 1 and 2). The 2D [^13^C, ^1^H] HSQC NMR spectra reveal that the TAR-G28U/C37A rescue mutation restores a GS-like conformation to >90% population, and a small residual ES2 population is possible given line broadening in the NMR spectra at residues that undergo exchange with ES2 (Fig 2C, Supplementary Fig 2A). These results highlight how mutations do not induce all-or-nothing behavior (*i.e.* GS or ES) but rather redistribute the ensemble along a continuous scale (Fig 1). Using 2D [^13^C, ^1^H] HSQC NMR, we validated the dominant conformation of all other ES2-stabilizing mutants and their corresponding rescue mutants and also assessed the degree to which they stabilize/destabilize the GS relative to the ES (Fig 2C, Supplementary Fig 2).

The mutants were tested in a Tat-dependent HIV-1 LTR-driven luciferase reporter assay (Fig 2D). If TAR-G28U forms the ES2 conformation within cells with an abundance comparable to that measured *in vitro*, and if the TAR ES2 conformation is indeed inactive, we would expect the mutant to show at least a 1000-fold reduction in activity relative to WT (an even greater level of inhibition is in principle possible if the exchange kinetics between the GS and ES are slow relative to protein binding). As positive controls, we first tested double mutants TAR-G36C/C29G and TAR-G28C/C37G, which adopt a predominantly GS-like conformation based on NMR (Fig 2C, Supplementary Fig 2C). As expected, the mutants did not have statistically different trans-activation activity relative to WT (two-way ANOVA, P > 0.05) (Fig 2D). In contrast, the trans-activation activity of the ES2-stabilizing TAR-G28U mutant fell below detectable levels with indistinguishable activity from the -Tat negative control (two-sided unpaired Student’s t-test, P > 0.05), corresponding to at least 100-fold decrease in activity (Fig 2D). The activity increased significantly (two-way ANOVA, P < 0.05) and was restored to WT TAR levels (two-way ANOVA, P > 0.05) upon introduction of a second rescue mutation (TAR-G28U/C37A) (Fig 2D). Although not statistically significant, the lower mean activity observed for TAR-G28U/C37A relative to WT could arise from incomplete restoration of the GS and/or because the sequence of the G28-C37 base pair is important for activity.

The ES2-stabilizing TAR-G28U mutant introduces a mismatch in the native GS conformation and we cannot rule out the possibility that the cellular environment does favor the GS but that the mismatch disrupts the ability of TAR to bind to its protein partners. Experiments were therefore repeated for the three other sets of mutants each targeting different TAR residues (TAR-A27C, TAR-G36U, and TAR-U38A) and the results were similar to those obtained for TAR-G28U (Fig 2D). All three ES2-stabilizing mutants drastically diminish TAR activity, while all but one GS-rescue mutation at least partially restores activity (two-way ANOVA, P < 0.05) (Fig 2D). For most cases, the effect of the mutation on activity *in vivo* correlated with the degree of ES-stabilization measured *in vitro*. For example, based on NMR, TAR-G36U and TAR-A27C bias the conformational equilibrium toward ES2 to a similar degree as TAR-G28U (Fig 2C), and correspondingly, their activities are indistinguishable from that observed for TAR-G28U (two-way Anova, P > 0.05) (Fig 2D). Based on NMR, TAR-U38A stabilizes ES2 to a smaller degree compared to other mutants (Fig 2C), and correspondingly it has marginally higher activity (∼6% of WT TAR) relative to other ES2-stabilizing mutants (Fig 2D). Likewise, the partial rescue of TAR-G36U/C29A in the transactivation assay mirrors the partial conformational rescue as assessed by NMR (Fig 2C and 2D), although we cannot rule out that the identity of the closing G36-C29 bp is also important for protein recognition (Fig 2C and 2D). Based on NMR, the rescue mutants TAR-U38A/A27U and TAR-A27C/U38G strongly stabilize the GS yet they only partially restore activity (Fig 2C and 2D). This not surprising given that in this case the mutations disrupt an U23•U38-A27 base triple known to be important for Tat binding(Puglisi et al., 1992, 1993).

ES1 is localized to the apical loop and does not change the TAR secondary structure as significantly as ES2. Therefore, it was expected that ES1 may not be an inactive conformation as could be assumed for ES2. Correspondingly, the two ES1-stabilizing mutants(Dethoff et al., 2012b) (TAR-C30U and TAR-A35G) did not significantly decrease trans-activation relative to WT activity (two-way ANOVA, P > 0.5) (Fig 2D).

Taken together, the agreement between the conformations determined *in vitro* by NMR and the cell-based trans-activation activity suggests that the energetic stability of ES2 relative to the GS are not substantially altered in the cell relative to the *in vitro* environment. The results also confirm that ES2 is inhibitory, as is expected given prior studies showing the importance of the TAR GS secondary structure for transactivation(Feng and Holland, 1988; Berkhout and Jeang, 1989; Dingwall et al., 1990; Roy et al., 1990; Churcher et al., 1993; Schulze-Gahmen et al., 2016).

To test the generality of our approach, we examined the relative energetic stability and abundance of ESs that have recently been characterized in HIV-1 RRE stem IIB. RRE RNA mediates nuclear export of incompletely spliced HIV-1 RNAs by cooperatively binding multiple copies of the viral Rev protein(Bartel et al., 1991). This assembly is initiated through binding of Rev to the purine rich region of RRE stem IIB(Bartel et al., 1991). Our recent work(Chu et al., 2019) showed that RRE stem IIB forms two non-native ESs (ES1 and ES2) both of which disrupt key structural elements recognized by Rev, including the G47-A73 and G48-G71 mismatches and U72 bulge (Fig 3A). ES1 reshuffles the lower bulge while ES2 remodels both the lower and upper bulge (Fig 3A)(Chu et al., 2019).

**Figure 3.**
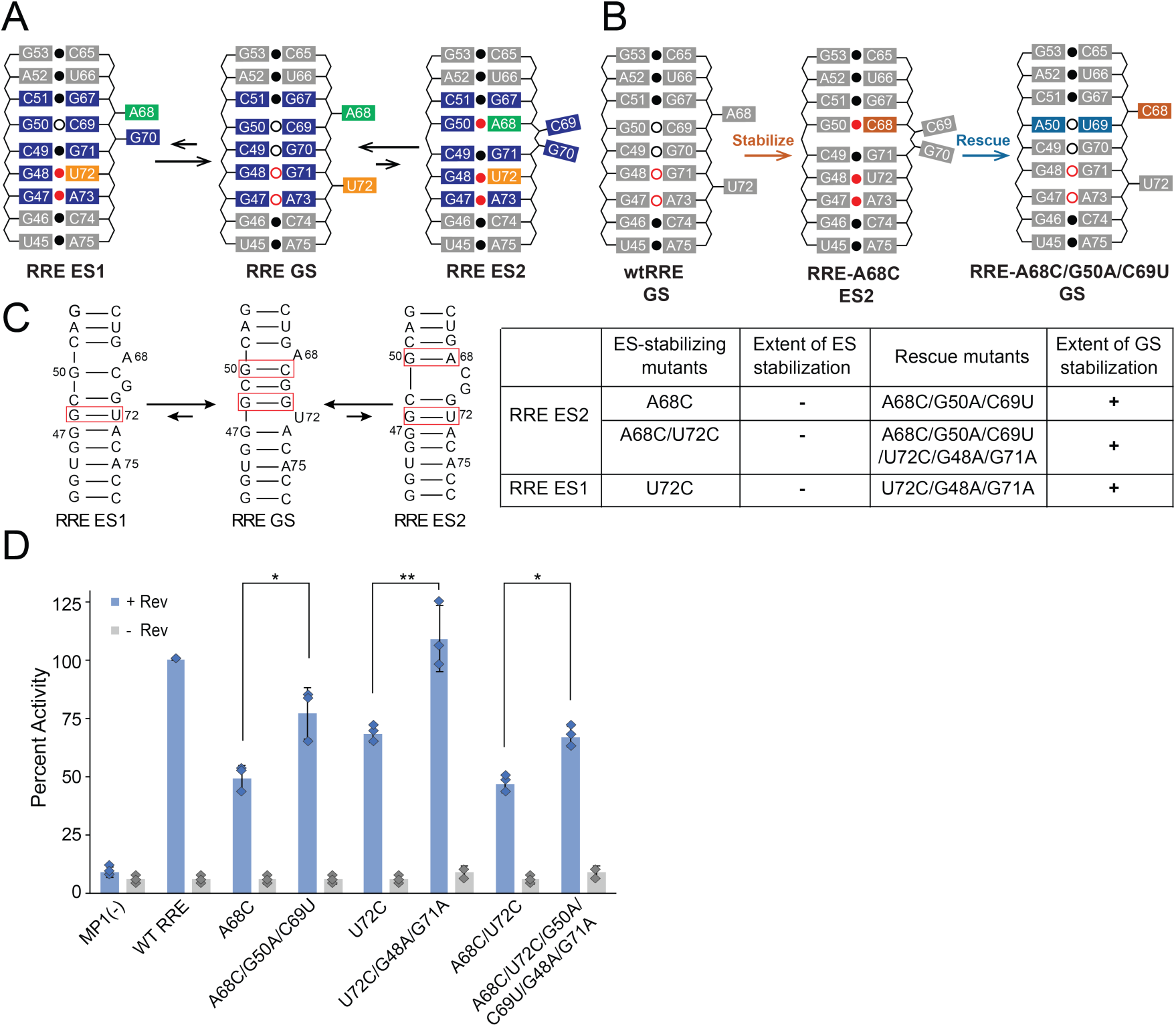
Probing the conformational equilibrium of HIV-1 RRE *in vitro* and *in vivo*. **A.** HIV-1 RRE RNA has two excited states that reshuffle the non-canonical base pairs at the purine-rich region (blue), U72 (yellow), and A68 (green) bulge. Weak base pairs are indicated with an open dot and non-Watson-Crick base pairs are indicated with a red dot. **B.** Example of the stabilize-and-rescue approach for trapping RRE ES2 using an ES-stabilizing mutant A68C (orange) and corresponding rescue mutations G50A and C69U (blue). Weak base pairs are indicated with an open dot and non-Watson-Crick base pairs are indicated with a red dot. **C.** Summary of all RRE mutants. Base pairs that were mutated are highlighted (red box). The extent of stabilization was estimated based on NMR line broadening (Supplementary Fig 6). (++) minimal line broadening, (+) partial line broadening, and (-) extensive line broadening. **D.** Rev-RRE dependent RNA export assays of RRE mutants. 293T cell were transfected with pFLuc-RRE reporter plasmid and an RLuc internal control in the presence (+ Rev) or absence (-Rev) of a Rev expression plasmid. At 48 hr post transfection, cell lysates were tested for luciferase activity. Reported values are the quotient of FLuc and RLuc activities with values normalized to WT RRE for every independent replicate reported as the mean ± SD. MP-1 is the control plasmid (no RRE). Statistical significance was determined after log-transformation of the data; *P < 0.05, **P < 0.01, ***P < 0.001 (two-sided ANOVA).

As was done for TAR, point substitution mutations stabilizing the RRE ESs and corresponding rescue mutations that restore the GS were designed and verified using NMR (Fig 3B and 3C, Supplementary note 1, Supplementary Figs 5-6). The mutants were then tested in an RRE-dependent RNA export assay to monitor RRE activity (Fig 3D). Due to the localized nature of the conformational change, it was challenging to design mutants that stabilize the RRE ESs to >99% population. Rather, extreme broadening of the NMR spectra indicate that RRE-A68C only partially stabilizes ES2 to a population ranging between 20% and 80% (Fig 3C and Supplementary Fig 6). This provides an important test for our approach because we expect to see smaller levels of attenuation of cellular activity as compared to TAR. The RRE stem IIB GS conformation is largely rescued by RRE-A68C/G50A/C69U based on the restoration of WT chemical shifts (Fig 3C and Supplementary Fig 6). Strikingly, as expected given that it only partially stabilizes the ES, RRE-A68C showed only a 2-fold reduction in RRE-mediated export relative to WT while the rescue mutant RRE-A68C/G50A/C69U restored activity to within ∼80% of WT (Fig 3D). Likewise, RRE-U72C partially stabilizes the cumulative ES1 + ES2 population to 20%-80% (Fig 3C), and correspondingly decreased export activity to ∼70% of WT, while the rescue mutant RRE-U72C/G48A/G71A restored activity to ∼100% of WT (Fig 3D). Another double mutant, RRE-A68C/U72C, which also partially stabilizes ES2 to 20%-80% based on NMR (Fig 3C), also shows a 2-fold reduction in RRE-mediated export to ∼50% of WT while the rescue mutant RRE-U72C/G48A/G71A/A68C/G50A/C69U restores activity to ∼70% of WT (Fig 3D). Similar results were also observed for Rev-RRE binding in cells using a previously described TAR-RRE hybrid trans-activation assay(Tiley et al., 1992) (Supplementary Fig 7), confirming that inhibition of RRE export is indeed due to inhibition of Rev-RRE binding.

Although the effects are smaller than those observed for TAR, the robust correlation between the impact on conformation and on RRE activity for different ES-stabilizing and rescue mutants indicates that stabilizing the RRE ES conformation is indeed inhibitory (Fig 3C, 3D and Supplementary Figs 5-7). Since the mutants only stabilize the RRE ES to ∼50% population, the 2-fold reduction in activity is consistent with the ES conformation being fully inhibitory and the observed remaining activity can be attributed to a residual GS population. Once again, if the cellular environment preferentially stabilizes the GS relative to the ES, we would have expected the mutations to have a smaller effect within cells than predicted from their equilibrium populations, which is not observed.

In conclusion, we have described a new approach to probe the conformational thermodynamics involving low-abundance short-lived RNA ESs within the functional cellular context. This approach can be generally applied to examine how the cellular environment impacts RNA secondary structural equilibria relative to *in vitro* conditions. The approach also provides a means by which to test the structure-dependence of RNA function within cells because it maximally changes RNA structure while minimally impacting sequence. Our results indicate that for TAR ES2 and RRE ES1 and ES2, the cellular environment does not substantially reduce the stability and abundance of the ES relative to the GS compared to the stability measured *in vitro*. Since none of the mutants exhibited lower biological activity than expected based on their *in vitro* stabilities, it is also unlikely that the cellular environment substantially stabilizes the ES relative to the GS compared to *in vitro* conditions. These findings validate an HIV therapeutic strategy that involves targeted stabilization of these inactive ES conformations.

## Acknowledgements

We would like to thank the Duke Magnetic Resonance Spectroscopy Center for NMR resources. This work was supported by the US National Institutes of Health (P50 GM103297 to B.R.C. and H.M.A.-H and F31 GM119306 to L.R.G).

## Author Contributions

The study was designed by L.R.G, C-C.C. and H.M.A.-H. Experiments were performed by L.R.G. C-C.C., H.P.B. and M.L.K. The manuscript was written by L.R.G., C-C.C., and H.M.A.-H. with input from all other co-authors.

## Methods

### RNA sample preparation

Unlabeled RNA samples (TAR-G28U, TAR-G28U/C37A, TAR-G36U, TAR-G36U/C29A, TAR-U38A, TAR-U38A/A27U, TAR-A27C, TAR-A27C/U38G, WT TAR, and uucgES2TAR) were prepared using a MerMade 6 DNA/RNA synthesizer (BioAutomation) for solid-phase oligonucleotide synthesis using standard phosphoramidite RNA chemistry and deprotection protocols. ^13^C/^15^N labeled RNA samples (RRE-A68C/G50A/C69U/U72C/G48A/G71A, RRE-A68C/U72C, RRE-U72C/G48A/G71A, RRE-U72C, RRE-A68C/G50A/C69U, RRE-A68C and TAR-G28U) used for NMR characterization were prepared by *in vitro* synthesis using T7 RNA polymerase (New England BioLabs), DNA template (Integrated DNA Technologies) containing the T7 promoter, and uniformly labeled ^13^C/^15^N-labeled nucleotide triphosphates (Cambridge Isotope Laboratories, Inc.). The transcription reaction was carried out at 37 °C for 12 hours and then filtered with a 0.2 mm filter.

All RNA samples were purified using a 20% (w/v) polyacrylamide gel with 8 M urea and 1 X Tris/borate/EDTA. The RNA was removed from the excised gel by electro-elution in 1X Tris/acetic acid/EDTA followed by ethanol precipitation. The RNA was annealed in water at a concentration of 50 mM by heating at 95 °C for 5 min followed by cooling on ice for 30 min. The RNA was then buffer exchanged using an Amicon Ultra-15 centrifugal filter (EMD Milipore) with a 3 KDa cutoff into NMR buffer [15 mM sodium phosphate, 25 mM sodium chloride, and 0.1 mM EDTA, pH 6.4]. Finally 10% (v/v) D_2_O was added to the sample before NMR data were collected. The final concentration of RNA in all samples was between 0.6 and 1.1 mM.

### NMR spectroscopy

#### Assessing impact of mutations using chemical shift mapping experiments

The impact of the mutations on RNA structure was initially assessed by recording 1D [^1^H] and 2D [^13^C-^1^H] SOFAST-HMQC NMR experiments(Sathyamoorthy et al., 2014) using unlabeled RNA samples. All NMR experiments were carried out at 298 K on a 600 MHz Bruker spectrometer equipped with HCN cryogenic probes. Data were processed using NMRpipe(Delaglio et al., 1995) and analyzed using SPARKY(Goddard and Kneller, n.d.).

#### TAR

Comparison of NMR spectra for each TAR mutant with corresponding spectra of WT TAR and a validated TAR ES2 mutant (UUCG-ES2 TAR)(Merriman et al., 2016; Clay et al., 2017) was used to assess the degree of ES-versus GS-stabilization. Although the entire spectrum was evaluated (Supplementary Fig 2), key residues used in this analysis include A22 C8-H8/C2-H2, U23 C6-H6, C24 C6-H6, A35 C8-H8, U40 C6-H6, and C41 C6-H6. To determine the degree of stabilization of ES-and GS-stabilizing mutants, the extent of broadening for these spectra was investigated. The broadening of TAR-U38A is apparent in many resonances, but particularly in the peak tentatively assigned to U23, which is not broadened out in the other ES2 traps. The broadening of TAR-G28U/C37A and TAR-G36U/C29A relative to the other rescue mutants is also particularly visible in U23.

#### RRE

Likewise, comparison of NMR spectra for each RRE mutant with corresponding spectra of WT RRE was used to assess the degree ES-versus GS-stabilization. The entire spectra were evaluated (Supplementary Fig 5), and the key resonances used in this analysis include A68 C8-H8, U72 C6-H6, G47 C8-H8, G48 C8-H8, and G71 C8-H8. The broadening of these resonances, which experience significant changes in chemical shift between the GS and ESs were used to estimate the relative stability of ESs versus GS.

#### ^15^N R_1ρ_ relaxation dispersion

^15^N R_1ρ_ relaxation dispersion (RD) NMR experiments were carried out using a 700 MHz Bruker Avance III spectrometer equipped with an HCN cryogenic probe. Measurements were carried out at 25°C on the 0.7 mM uniformly ^13^C/^15^N-labeled TAR-G28U sample. R_1ρ_ RD for N3 spins were measured using a 1D acquisition scheme(Nikolova et al., 2012) using spinlock powers (w2p^-1^ Hz) and offsets (W2p^-1^ Hz) listed in Supplementary Table 1 and using 7 time delay points between 0 and 120 ms. For each delay time point, peak intensities were obtained using NMRPipe and R_1ρ_ values were calculated by fitting to a monoexponential decay function using an in-house python script(Kimsey et al., 2015). Errors in R_1ρ_ were estimated using Monte Carlo simulations with 500 iterations as previously described(Bothe et al., 2014). The RD data was analyzed to obtain exchange parameters through numerical solutions of the Bloch-McConnell equations(Korzhnev et al., 2005) using an in-house python script(Kimsey et al., 2018). As expected the TAR-U42-N3 negative control showed no dispersion. The TAR-U38-N3 data was fit with an individual two-state model using ground state effective field alignment. Model selection was preformed as previously described(Kimsey et al., 2018) using the Akaike’s (*w*AIC) and Bayesian information criterion (*w*BIC) weights to select the model with the highest relative probability(Burnham and Anderson, 2004; Wagenmakers and Farrell, 2004).

### Plasmid construction

The Tat and Rev expression plasmids, pcTat(Tiley et al., 1992) and pcRev(Malim et al., 1988), pcTat/Rev(Tiley et al., 1992), and pBC12-CMV(Tiley et al., 1992) have been described previously. The Renilla luciferase (RLuc) internal control was generated by PCR amplification of the RLuc ORF, which was then cloned into pcDNA3 (Invitrogen) using unique HindIII and XhoI restriction sites. The Firefly luciferase (FLuc) reporter plasmids with TAR and RRE, both wild-type and mutants, were constructed as follows. The pFLuc-TAR plasmid was generated by cloning the FLuc gene into pcDNA3 (Invitrogen) using NotI and XhoI restriction sites to generate pcDNA-FLuc. The HIV-LTR was next cloned as a Bam HI/Hind III fragment into pcDNA-FLuc digested with Bgl II and Hind III, which removes the entire CMV promoter. TAR mutants were introduced at Bgl II and Hind III restriction sites using annealed DNA oligos (IDT) that were designed to have overhangs complimentary to Bgl II and Hind III. The pFLuc-TAR/RRE plasmid was generated with the same Bgl II and Hind III restriction sites using annealed DNA oligos (IDT) containing RRE stem II sequence (66nt). The pFLuc-RRE was derived from pDM128-PL(Fridell et al., 1993). The CAT gene was replaced with the FLuc gene using Sal I and Bgl II restriction sites (pMP-1). The RRE and variants were generated by ligating in the whole RRE (∼350 nt) inserts amplified using two-step recombinant PCR, using the HIV-1 proviral clone pNL4-3 as the template, (AIDS reagent cat #114). The PCR products of RRE and variants were then inserted to pMP-1 at Kpn I/Xba I restriction sites.

### Trans-activation assays

HeLa cells were maintained in Dulbecco’s modified Eagle medium (DMEM) supplemented with 10% fetal bovine serum (FBS) and 0.1% gentamycin at 37 °C and 5% CO_2_. Cells were plated to 3 × 10^5^ cells per well in 12-well plates and transfected using PEI with 250 ng pFLuc-TAR reporter plasmid, 20 ng RLuc internal control, 20 ng pcTat, and pBC12-CMV filler DNA up to 1750 ng total DNA. Media was replaced at 24 h post-transfection and cells were lysed at 48 h post-transfection using 200 ml passive lysis buffer (Promega) followed by freezing at −80 °C. FLuc and RLuc activity was measured using a dual-luciferase reporter assay system (Promega). Reported values are the quotient of FLuc activity and RLuc activity with values normalized to WT TAR for every replicate reported as the mean ± SD (Fig 2D).

### Rev-RRE binding assay

For Rev-RRE binding assay, HeLa cells (2 × 10^5^ cells per well) were transfected using PEI with 350 ng pFLuc-TAR/RRE reporter plasmid, 20 ng RLuc internal control, and 100 ng pcTat/Rev or 100 ng pcTat, and pBC12-CMV filler DNA up to 1370 ng total DNA. The cell lysate preparation and luciferase measurement are as described above for the trans-activation assay. Reported values are the quotient of FLuc activity and RLuc activity with values normalized to WT RREII for every replicate reported as the mean ± SD (Supplementary Fig 7).

### RRE-mediated RNA export assay

293T cells were maintained in DMEM supplemented with 10% FBS and 0.1% gentamycin at 37 °C and 5% CO_2_. Cells were plated to 2 × 10^5^ cells per well in 12-well plates and transfected using PEI with 5 ng pFLuc-RRE reporter plasmid, 5 ng RLuc internal control, 1 ng pcRev, and pBC12-CMV filler DNA up to 1010 ng total DNA. Media was replaced at 24 h post-transfection and cells were lysed at 48 h post-transfection using 200 ml passive lysis buffer (Promega). FLuc and RLuc activity was measured using a dual-luciferase reporter assay system (Promega). Reported values are the quotient of FLuc activity and RLuc activity with values normalized to WT RREII activity for every replicate reported as the mean ± SD (Fig 3D).

### Statistical analysis

Statistical analysis was done using the program JMP (JMP Pro, Version 14, SAS Institute Inc., Cary, NC, 1989-2019). The data was not normalized to WT TAR for the trans-activation assay because there was not a significant systematic effect of replicate on the data. For both RRE assays, the data was normalized to WT RREII because there was a significant systematic effect of replicate on the data. For all three assays, the FLuc/RLuc quotient was log transformed for statistical analysis. A two-sided unpaired Student’s t-test was used to compare the +Tat and – Tat distributions for the trans-activation assays and a two-sided ANOVA was used to compare the difference between distributions (+Tat and –Tat; +Rev and –Rev; or +Tat/Rev and +Tat) for different samples for all three assays.

## References

Abeysirigunawardena, S.C., Kim, H., Lai, J., Ragunathan, K., Rappé, M.C., Luthey-schulten, Z., Ha, T., Woodson, S.A. (2017). Evolution of protein-coupled RNA dynamics during hierarchical assembly of ribosomal complexes. Nature Commun. 8, 492.

Bartel, D.P., Zapp, M.L., Green, M.R., Szostak, J.W. (1991). HIV-1 Rev Regulation Involves Recognition of Non-Watson-Crick Base Pairs in Viral RNA. Cell 67, 529–536.

Beaudoin, J.D., Novoa, E.M., Vejnar, C.E., Yartseva, V., Takacs, C.M., Kellis, M., Giraldez, A.J. (2018). Analyses of mRNA structure dynamics identify embryonic gene regulatory programs. Nature Struct Mol Biol 25, 677–686.

Berkhout, B., Jeang, K.T. (1989). trans activation of human immunodeficiency virus type 1 is sequence specific for both the single-stranded bulge and loop of the trans-acting-responsive hairpin: a quantitative analysis. J Virol 63, 5501–5504.

Bieniasz, P.D., Grdina, T.A., Bogerd, H.P., Cullen, B.R. (1999). Recruitment of cyclin T1/P-TEFb to an HIV type 1 long terminal repeat promoter proximal RNA target is both necessary and sufficient for full activation of transcription. PNAS 96, 7791–7796.

Bothe, J.R., Stein, Z.W., Al-Hashimi, H.M. (2014). Evaluating the uncertainty in exchange parameters determined from off-resonance R1ρ relaxation dispersion for systems in fast exchange. J of Magn Reson 244, 18–29.

Breaker, R.R. (2011). Prospects for riboswitch discovery and analysis. Mol Cell 43, 867–879.

Burnham, K.P., Anderson, D.R. (2004). Multimodel inference: Understanding AIC and BIC in model selection. Sociological Methods & Research 33, 261–304.

Rangadurai, A., Szymaski, E.S., Kimsey, I.J., Shi, H., Al-Hashimi, H.M. (2019). Characterizing micro-to-millisecond chemical exchange in nucleic acids using off-resonance R1ρ relaxation dispersion. Prog Nucl Magn Reson Spectrosc 112, 55–102.

Chavali, S.S., Bonn-Breach, R., Wedekind, J.E. (2019). Face-time with TAR: Portraits of an HIV-1 RNA with diverse modes of effector recognition relevant for drug discovery. J Biol Chem 294, 9326–9341.

Chu, C.C., Plangger, R., Kreutz, C., Al-Hashimi, H.M. (2019). Dynamic ensemble of HIV-1 RRE stem IIB reveals non-native conformations that disrupt the Rev-binding site. Nucleic Acids Res 47, 7105–7117.

Churcher, M.J., Lamont, C., Hamy, F., Dingwall, C., Green, S.M., Lowe, A.D., Butler, J.G., Gait, M.J., Karn, J. (1993). High affinity binding of TAR RNA by the human immunodeficiency virus type-1 tat protein requires base-pairs in the RNA stem and amino acid residues flanking the basic region. J Mol Biol 230, 90–110.

Clay, M.C., Ganser, L.R., Merriman, D.K., Al-Hashimi, H.M. (2017). Resolving sugar puckers in RNA excited states exposes slow modes of repuckering dynamics. Nucleic Acids Res 45, e134.

Connelly, C.M., Moon, M.H., Schneekloth, J.S. Jr (2016). The emerging role of RNA as a therapeutic target for small molecules. Cell Chem Biol 23, 1077–1090.

Cruz, J.A., Westhof, E. (2009). The dynamic landscapes of RNA architecture. Cell 136, 604–609.

Delaglio, F., Grzesiek, S., Vuister, G.W., Zhu, G., Pfeifer, J., Bax, A., 1995. NMRPipe: a multidimensional spectral processing system based on UNIX pipes. J Biomol NMR 6, 277–293.

Dethoff, E.A., Chugh, J., Mustoe, A.M., Al-Hashimi, H.M. (2012a). Functional complexity and regulation through RNA dynamics. Nature 482, 322–330.

Dethoff, E.A., Petzold, K., Chugh, J., Casiano-Negroni, A., Al-Hashimi, H.M. (2012b). Visualizing transient low-populated structures of RNA. Nature 491, 724–728.

Dibrov, S.M., Parsons, J., Carnevali, M., Zhou, S., Rynearson, K.D., Ding, K., Garcia Sega, E., Brunn, N.D., Boerneke, M.A., Castaldi, M.P., Hermann, T. (2014). Hepatitis C virus translation inhibitors targeting the internal ribosomal entry site. J Med Chem 57, 1694–1707.

Dingwall, C., Ernberg, I., Gait, M.J., Green, S.M., Heaphy, S., Karn, J., Lowe, A.D., Singh, M., Skinner, M.A. (1990). HIV-1 tat protein stimulates transcription by binding to a U-rich bulge in the stem of the TAR RNA structure. EMBO J 9, 4145–4153.

Feng, S., Holland, E.C. (1988). HIV-1 tat trans-activation requires the loop sequene within tar. Nature 334, 165–167.

Fridell, R.A., Partin, K.M., Carpenter, S., Cullen, B.R. (1993). Identification of the Activation Domain of Equine Infectious Anemia Virus Rev. J Virol 67, 7317–7323.

Fujinaga, K., Irwin, D., Huang, Y., Taube, R., Kurosu, T., Peterlin, B.M. (2004). Dynamics of human immunodeficiency virus transcription: P-TEFb phosphorylates RD and dissociates negative effectors from the transactivation response element. Mol Cell Biol 24, 787–795.

Fürtig, B., Nozinovic, S., Reining, A., Schwalbe, H. (2015). Multiple conformational states of riboswitches fine-tune gene regulation. Curr Opin Struct Biol 30, 112–124.

Ganser, L.R., Kelly, M.L., Herschlag, D., Al-Hashimi, H.M. (2019). The roles of structural dynamics in the cellular functions of RNAs. Nat Rev Mol Cell Biol 20, 474–489.

Ganser, L.R., Lee, J., Rangadurai, A., Merriman, D.K., Kelly, M.L., Kansal, A.D., Sathyamoorthy, B., Al-hashimi, H.M. (2018). High-performance virtual screening by targeting a high-resolution RNA dynamic ensemble. Nat Struct Mol Biol 25, 425–434.

Goddard, T.D., Kneller, D.G., SPARKY 3, University of California, San Francisco

Guo, J.U., Bartel, D.P. (2016). RNA G-quadruplexes are globally unfolded in eukaryotic cells and depleted in bacteria. Science 353, aaf5371.

He, N., Liu, M., Hsu, J., Xue, Y., Chou, S., Burlingame, A., Krogan, N.J., Alber, T., Zhou, Q. (2010). HIV-1 Tat and host AFF4 recruit two transcription elongation factors into a bifunctional complex for coordinated activation of HIV-1 transcription. Mol Cell 38, 428–438.

Helmling, C., Klötzner, D., Sochor, F., Mooney, R.A., Wacker, A., Landick, R., Fürtig, B., Heckel, A., Schwalbe, H. (2018) Life times of metastable states guide regulatory signaling in transcriptional riboswitches. Nat Commun 9, 944.

Hermann, T. (2002). Rational ligand design for RNA: The role of static structure and conformational flexibility in target recognition. Biochimie 84, 869–875.

Ivanov, D., Kwak, Y.T., Guo, J., Gaynor, R.B. (2000). Domains in the SPT5 protein that modulate its transcriptional regulatory properties. Mol Cell Biol 20, 2970–2983.

Kim, J.N., Breaker, R.R. (2008). Purine sensing by riboswitches. Biol Cell 100, 1–11.

Kim, Y.K., Bourgeois, C.F., Isel, C., Churcher, M.J., Karn, J. (2002). Phosphorylation of the RNA polymerase II carboxyl-terminal domain by CDK9 is directly responsible for human immunodeficiency virus type 1 Tat-activated transcriptional elongation. Mol Cell Biol 22, 4622–4637.

Kimsey, I.J., Petzold, K., Sathyamoorthy, B., Stein, Z.W., Al-hashimi, H.M. (2015). Visualizing transient Watson-Crick-like mispairs in DNA and RNA duplexes. Nature 519, 315–320.

Kimsey, I.J., Szymanski, E.S., Zahurancik, W.J., Shakya, A., Xue, Y., Chu, C.C., Sathyamoorthy, B., Suo, Z., Al-Hashimi, H.M. (2018). Dynamic basis for dG • dT misincorporation via tautomerization and ionization. Nature 554, 195–201.

Korzhnev, D.M., Orekhov, V.Y., Kay, L.E. (2005). Off-resonance R(1rho) NMR studies of exchange dynamics in proteins with low spin-lock fields: an application to a Fyn SH3 domain. J Am Chem Soc 127, 713–21.

Lee, J., Dethoff, E.A., Al-Hashimi, H.M. (2014). Invisible RNA state dynamically couples distant motifs. Proc Natl Acad Sci 111, 9485–9490.

Leulliot, N., Varani, G. (2001). Current Topics in RNA-Protein Recognition: Control of Specificity and Biological Function through Induced Fit and Conformational Capture. Biochemistry 40, 7947–7956.

Li, H., Aviran, S. (2018). Statistical modeling of RNA structure profiling experiments enables parsimonious reconstruction of structure landscapes. Nat Commun 9, 606.

Malim, M.H., Hauber, J., Fenrick, R., Cullen, B.R. (1988). Immunodeficiency virus rev trans-activator modulates the expression of the viral regulatory genes. Nature 335, 181–183.

Merriman, D.K., Xue, Y., Yang, S., Kimsey, I.J., Shakya, A., Clay, M., Al-Hashimi, H.M. (2016). Shortening the HIV-1 TAR RNA bulge by a single nucleotide preserves motional modes over a broad range of time scales. Biochemistry 55, 4445–4456.

Mustoe, A.M., Brooks, C.L., Al-Hashimi, H.M. (2014). Hierarchy of RNA functional dynamics. Annu Rev Biochem 83, 441–466.

Mustoe, A.M., Busan, S., Rice, G.M., Hajdin, C.E., Peterson, B.K., Ruda, V.M., Kubica, N., Nutiu, R., Baryza, J.L., Weeks, K.M. (2018). Pervasive Regulatory Functions of mRNA Structure Revealed by High-Resolution SHAPE Probing. Cell 173, 181–195.

Nichols, P.J., Henen, M.A., Born, A., Strotz, D., Güntert, P., Vögeli, B. (2018). High-resolution small RNA structures from exact nuclear Overhauser enhancement measurements without additional restraints. Commun Biol 1, 61.

Nikolova, E.N., Gottardo, F.L., Al-Hashimi, H.M. (2012). Probing transient Hoogsteen hydrogen bonds in canonical duplex DNA using NMR relaxation dispersion and single-atom substitution. J Am Chem Soc 134, 3667–3670.

Pham, V. V, Salguero, C., Khan, S.N., Meagher, J.L., Brown, W.C., Humbert, N., de Rocquigny, H., Smith, J.L., D’Souza, V.M. (2018). HIV-1 Tat interactions with cellular 7SK and viral TAR RNAs identifies dual structural mimicry. Nat commun 9, 4266.

Puglisi, J.D., Chen, L., Blanchard, S., Frankel, A.D. (1995). Solution structure of a bovine immunodeficiency virus tat-TAR peptide-RNA complex. Science 270, 1200–1203.

Puglisi, J.D., Chen, L., Frankel, A.D., Williamson, J.R. (1993). Role of RNA structure in arginine recognition of TAR RNA. Proc Natl Acad Sci 90, 3680–3684.

Puglisi, J.D., Tan, R., Calnan, B.J., Frankel, A.D., Williamson, J.R. (1992). Conformation of the TAR RNA-arginine complex by NMR spectroscopy. Science 257, 76–80.

Rogers, E., Heitsch, C.E. (2014). Profiling small RNA reveals multimodal substructural signals in a Boltzmann ensemble. Nucleic Acids Res 42, e171.

Roy, S., Delling, U., Chen, C.H., Rosen, C.A., Sonenberg, N. (1990). A bulge structure in HIV-1 TAR RNA is required for Tat binding and Tat-mediated trans-activation. Genes Dev 4, 1365–73.

Salmon, L., Yang, S., Al-Hashimi, H.M. (2014). Advances in the determination of nucleic acid conformational ensembles. Annu Rev Phys Chem 65, 293–316.

Sathyamoorthy, B., Lee, J., Kimsey, I., Ganser, L.R., Al-Hashimi, H. (2014). Development and application of aromatic [13C, 1H] SOFAST-HMQC NMR experiment for nucleic acids. J Biomol NMR 60, 77–83.

Schroeder, S.J. (2018). Challenges and approaches to predicting RNA with multiple functional structures. RNA 24, 1615–1624.

Schulze-Gahmen, U., Echeverria, I., Stjepanovic, G., Bai, Y., Lu, H., Schneidman-Duhovny, D., Doudna, J.A., Zhou, Q., Sali, A., Hurley, J.H. (2016). Insights into HIV-1 proviral transcription from integrative structure and dynamics of the Tat:AFF4:P-TEFb:TAR complex. Elife 5, e15910.

Schulze-gahmen, U., Hurley, J.H. (2018). Structural mechanism for HIV-1 TAR loop recognition by Tat and the super elongation complex. Proc Natl Acad Sci 115, 12973–12978.

Sekhar, A., Kay, L.E. (2013). NMR paves the way for atomic level descriptions of sparsely populated, transiently formed biomolecular conformers. Proc Natl Acad Sci 110, 12867–12874.

Shi, X., Huang, L., Lilley, D.M., Harbury, P.B., Herschlag, D. (2016). The solution structural ensembles of RNA kink-turn motifs and their protein complexes. Nat Chem Biol 12, 146–152.

Skrynnikov, N.R., Dahlquist, F.W., Kay, L.E. (2002). Reconstructing NMR spectra of “invisible” excited protein states using HSQC and HMQC experiments. J Am Chem Soc 124, 12352–12360.

Sobhian, B., Laguette, N., Yatim, A., Nakamura, M., Levy, Y., Kiernan, R., Benkirane, M. (2010). HIV-1 Tat assembles a multifunctional transcription elongation complex and stably associates with the 7SK snRNP. Mol Cell 38, 439–451.

Spasic, A., Assmann, S.M., Bevilacqua, P.C., Mathews, D.H. (2018). Modeling RNA secondary structure folding ensembles using SHAPE mapping data. Nucleic Acids Res 46, 314–323.

Stelzer, A.C., Frank, A.T., Kratz, J.D., Swanson, M.D., Gonzalez-Hernandez, M.J., Lee, J., Andricioaei, I., Markovitz, D.M., Al-Hashimi, H.M. (2011). Discovery of selective bioactive small molecules by targeting an RNA dynamic ensemble. Nat Chem Biol 7, 553–559.

Sun, L., Fazal, F.M., Li, P., Broughton, J.P., Lee, B., Tang, L., Huang, W., Kool, E.T., Change, H.Y., Zhang, Q.C. (2019). RNA structure maps across mammalian cellular compartments. Nat Struct Mol Biol 26, 322–330.

Tian, S., Das, R. (2016). RNA structure through multidimensional chemical mapping. Q Rev Biophys 49, e7.

Tiley, L.S., Madore, S.J., Malim, M.H., Cullen, B.R. (1992). The VP16 transcription activation domain is functional when targeted to a promoter-proximal RNA sequence. Genes Dev 6, 2077–2087.

Wagenmakers, E.J., Farrell, S. (2004). AIC model selection using Akaike weights. Psychon Bull Rev 11, 192–196.

Walter, F., Vicens, Q., Westhof, E. (1999). Aminoglycoside – RNA interactions. Curr Opin Chem Biol 3, 694–704.

Wei, P., Garber, M.E., Fang, S.M., Fischer, W.H., Jones, K.A. (1998). A novel CDK9-associated C-type cyclin interacts directly with HIV-1 Tat and mediates its high-affinity, loop-specific binding to TAR RNA. Cell 92, 451–462.

Williamson, J.R. (2000). Induced fit in RNA – protein recognition. Nat Struct Biol 7, 834–837.

Woods, C.T., Lackey, L., Williams, B., Dokholyan, N.V, Gotz, D., Laederach, A. (2017). Comparative Visualization of the RNA Suboptimal Conformational Ensemble In Vivo. Biophys J 113, 290–301.

Xue, Y., Gracia, B., Herschlag, D., Russell, R., Al-hashimi, H.M. (2016). Visualizing the formation of an RNA folding intermediate through a fast highly modular secondary structure switch. Nat Commun 7, 11768.

Xue, Y., Kellogg, D., Kimsey, I.J., Sathyamoorthy, B., Stein, Z.W., McBrairty, M., Al-Hashimi, H.M. (2015). Characterizing RNA excited states using NMR relaxation dispersion. In Methods in Enzymology, Elsevier Inc., pp. 39–73.

Zhao, B., Guffy, S.L., Williams, B., Zhang, Q. (2017). An excited state underlies gene regulation of a transcriptional riboswitch. Nat Chem Biol 13, 968–974.

Zhuang, X., Bartley, L.E., Babcock, H.P., Russell, R., Ha, T., Herschlag, D., Chu, S. (2000). A Single-Molecule Study of RNA Catalysis and Folding. Science 288, 2048–2052.

Zhuang, X., Kim, H., Pereira, M.J., Babcock, H.P., Walter, N.G., Chu, S. (2002). Correlating Structural Dynamics and Function in Single Ribozyme Molecules. Science 296, 1473–1477.

