## Supplementary Information for "Probing RNA conformational equilibria within the functional cellular context"

**Supplementary Note 1: Mutant design and validation**

*TAR*

Prior studies showed that TAR excited state (ES) conformations can be stabilized relative to the GS with TAR-A35C or TAR-C30U point substitution mutations in the case of ES1 and TAR-G28U in the case of ES2[1,2]. To test the robustness of our results and to control for minor structural effects of the mutations, we used the secondary structure prediction program MC-Fold[3] to guide the design of additional ES2 stabilizing mutants, TAR-U38A, TAR-A27C, TAR-G36U, TAR-U25A, TAR-C24G and TAR-C39G (Supplementary Fig 1a-c). Based on structure prediction[3], all mutants were predicted to adopt ES2 either as the most favored conformation or just marginally less stable than a very similar conformation with a C24 bulge rather than the U23 bulge (Supplementary Fig 1a-c). NMR chemical shift fingerprinting was used to assess the dominant structure and the extent of stabilization of each of these mutants (see methods, Supplementary Fig 2a-b). Mutants TAR-G28U, TAR-U38A, TAR-A27C, and TAR-G36U were shown to favor the ES2-conformation over the ground state (GS) conformation (Supplementary Fig 2a). Each of these ES2-stabilizing mutants replaces a mispair in the ES with Watson-Crick (WC) base pair (bp), while simultaneously replacing a WC bp in the GS conformation with a mispair. Conversely, mutants TAR-U25A and TAR-C24G, which replace a mispair in the ES with a WC without destabilizing the GS, were shown to maintain the GS conformation (Supplementary Fig 2b). Likewise, C39G, which switches a junctional mispair to a WC bp in the ES and switches a junctional WC bp to a mispair in the GS, causes significant broadening of the NMR spectra for residues in and around the bulge, but maintains the GS conformation overall (Supplementary Fig 2b). Rescue mutants were designed for all validated TAR ES2-stabilizing mutants that restore the WC bp in the GS and introduce a new mispair to the ES. Structure prediction with MC-Fold did not necessarily predict each rescue to favor the GS conformation (Supplementary Fig 1d), but NMR chemical shift mapping suggests that they do (Supplementary Fig 2a). As additional controls, two mutants were designed that are expected to destabilize the ESs relative to the GS by inverting a WC bp in the GS (Supplementary Fig 1e). These mutants were verified to favor the GS using NMR chemical shift fingerprinting (Supplementary Fig 2c). Together, these results highlight the importance of validating the conformational effects of mutants with NMR.

A previous study found that the ES2-stabilizing mutant, TAR-G28U, experiences a small amount of dimerization at NMR concentrations and two imino peaks were assigned to the inter-molecular U31-G32 base pair[4]. Evaluation of the other TAR mutants using 1D NMR of the imino protons reveal that the other ES2-stabilizing mutants and, to a lesser extent, rescue mutants also have minor peaks corresponding to dimerization (Supplementary Fig 3a,b). The TAR GS-stabilizing control mutants do not have peaks corresponding to dimerization (Supplementary Fig 3c), supporting the conclusion that the ES2 structure promotes dimerization. The dimer peaks are largest for TAR-U38A, which may, at least in part, explain the observed increased broadening in its 2D aromatic spectrum.

*RRE*

Our recent study showed that RRE excited state (ES) conformations can be stabilized relative to the GS with A68C and U72C point substitution mutations in the case of ES2 and ES1+ES2, respectively[5]. These mutations target nucleotides that do not play essential roles in RRE recognition and were predicted to fold predominantly into the ES conformation using MC-fold (Supplementary Fig 5a-b) [3]. To test robustness of results, an additional ES2-stabilizing mutant was designed that involves two point substitution mutations RRE-A68C/U72C which are predicted to stabilize the ES2 (Supplementary Fig 5b). Using NMR line broadening analysis, all three ES-stabilizing mutants were shown to stabilize the ESs to a population ranging between 20% and 80% based on the broadening of resonances that undergo chemical shift changes between the GS and the ES (Supplementary Fig 6). Recues for all RRE ES-stabilizing mutants were also designed here to demonstrate the structure dependency to RRE activity (Supplementary Fig 5c). The RRE-A68C rescue mutant (RRE-

A68C/G50A/C69U) was shown to stabilize the GS based on the overall agreement with the wtRREII spectra as well as the restoration of the U72 resonance, which is broadened out in the RRE-A68C ES2-stabilizing mutant. Similarly, the RRE-U72C rescue mutant (RRE- U72C/G48A/G71A) was also shown to stabilize the GS based on the overall agreement with the wtRREII spectra as well as the restoration of the A68 resonance, which is broadened out in the RRE-U72C ES2-stabilizing mutant. The RRE-A68C/U72C rescue mutant, RRE- A68C/G50A/C69U/U72C/G48A/G71A, six point substitutions were introduced to restore the GS conformation which was confirmed based on the overall agreement of its NMR spectra to that of wtRREII.

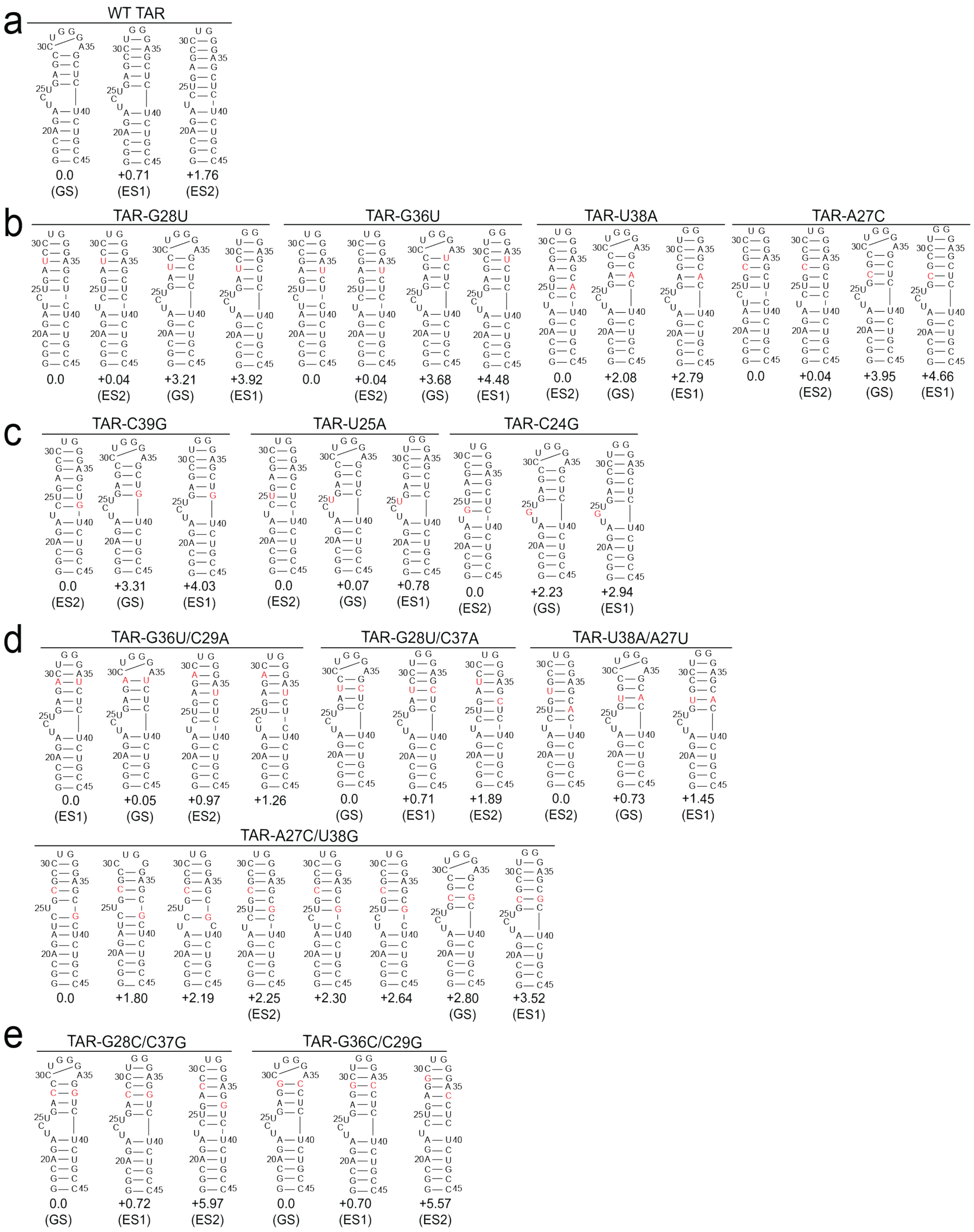


**Supplementary Figure 1**. MC-Fold predicted energies (kcal/mol) relative to the GS energy for the GS, ES1, and ES2 conformations of each TAR mutant as well as any conformations with lower predicted energy than the conformation that is being stabilized. Results shown for **a.** wtTAR, **b.** ES2-stabilizing mutants, **c.** Predicted ES2-stabilizing mutants that did not stabilize ES2 based on NMR, **d.** Rescue mutants, and **e.** GS-stabilizing controls.

a

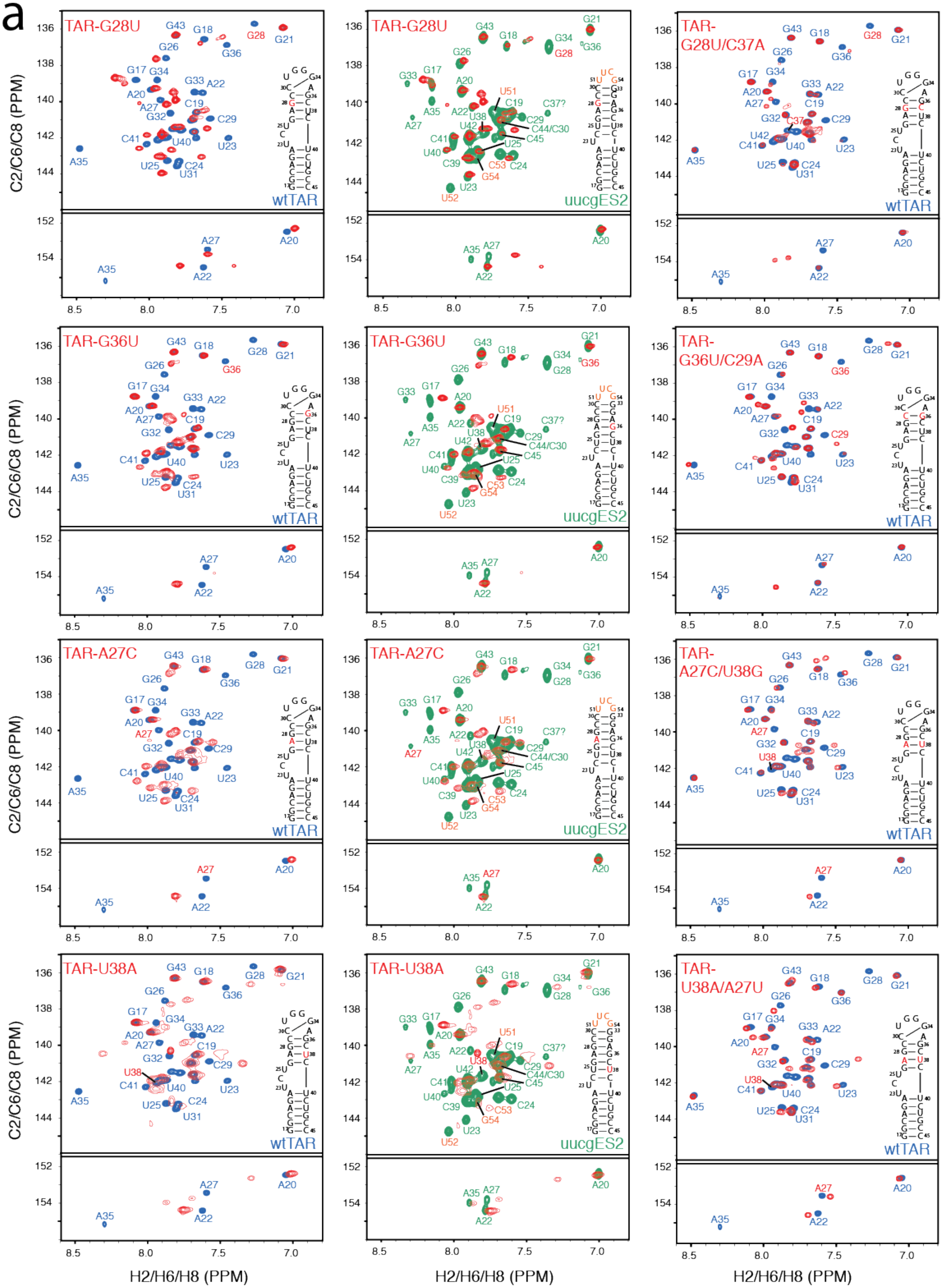

TAR-G28U
wtTAR
uucgES2
TAR-G28U/C37A
TAR-G36U
TAR-G36U/C29A
TAR-A27C
TAR-A27C/U38G
TAR-U38A
TAR-U38A/A27U
C2/C6/C8 (PPM)
H2/H6/H8 (PPM)

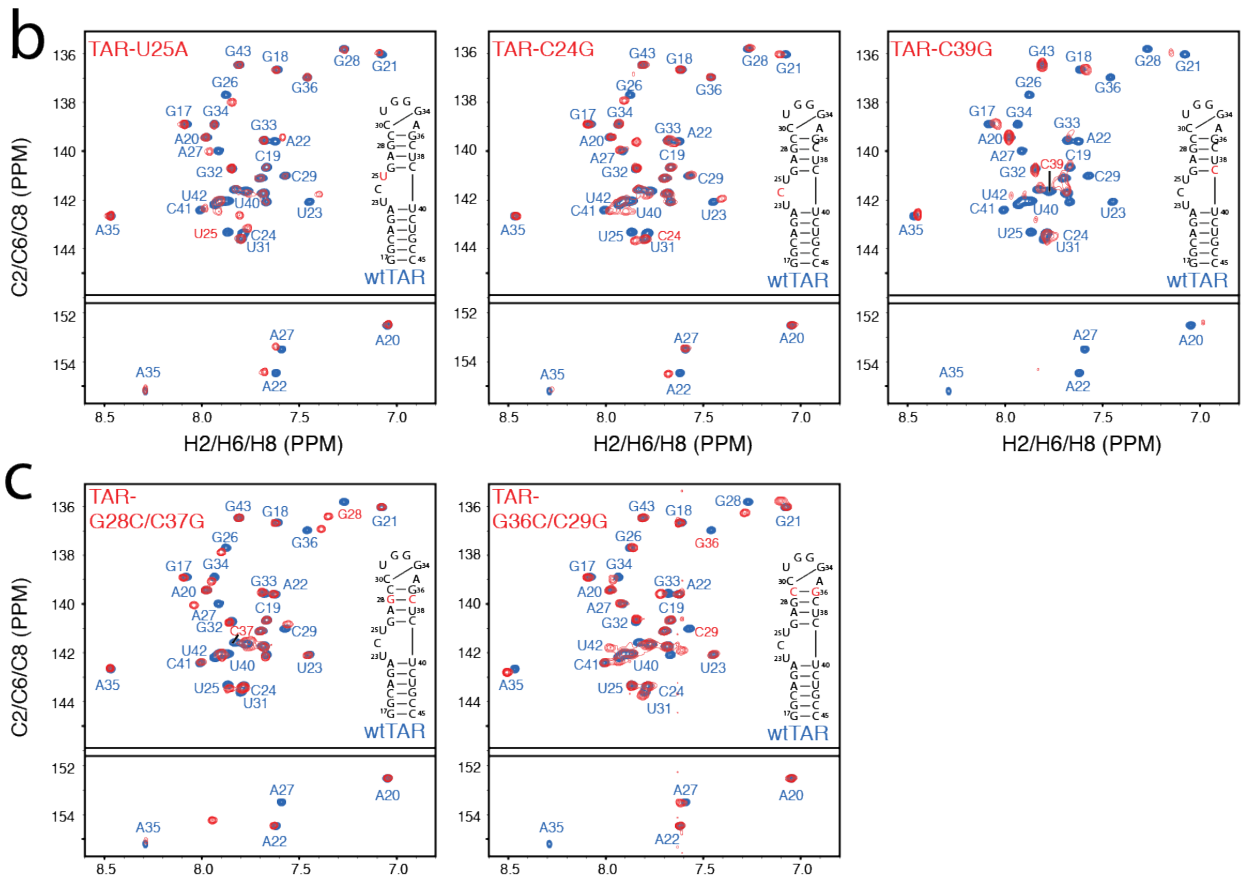


**Supplementary Figure 2.** NMR chemical shift mapping of TAR mutants (red) compared to wtTAR (blue) or a well-validated ES2-stabilizing mutant, UUCG-ES2 TAR (green) for **a.** ES2-stabilizing mutants and their respective rescue mutants, **b.** Predicted ES2-stabilizing mutants that maintain the GS-conformation, and **c.** GS-stabilizing control mutants.

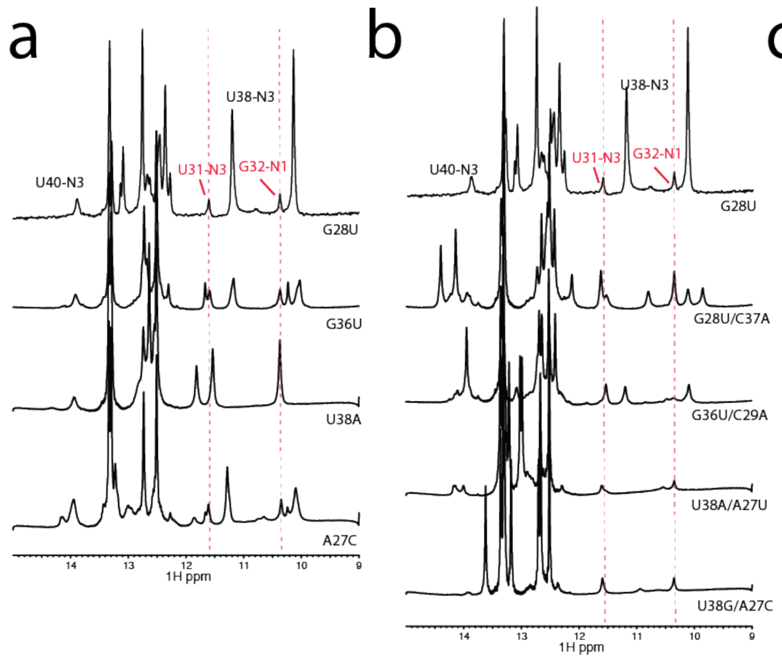


**Supplementary Figure 3.** 1D $^{1}$H SOFAST-HMQC NMR experiments on TAR mutants for **a.** ES2-stabilizing mutants, **b.** rescue mutants, and **c.** GS-stabilizing mutants. Red dotted lines indicate the chemical shifts of the peaks corresponding to dimerization based on the TAR-G28U mutant.

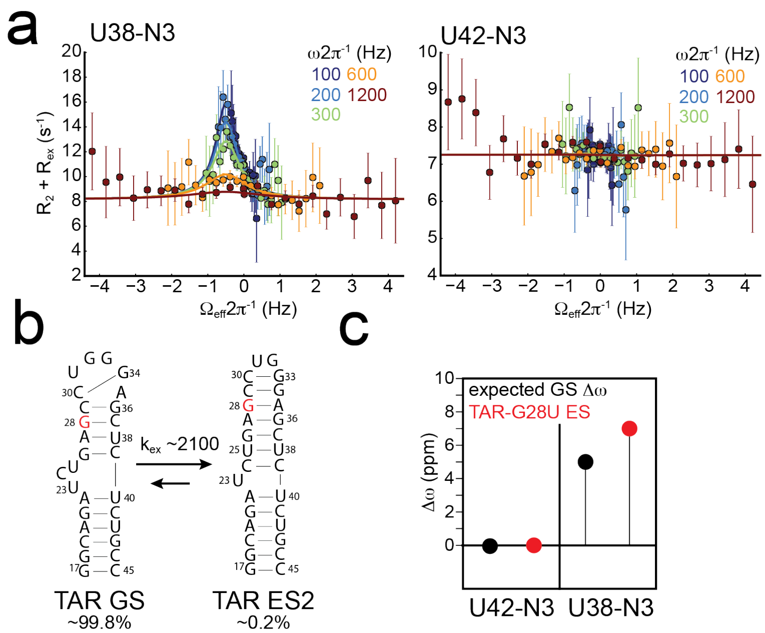


**Supplementary Figure 4:** NMR relaxation dispersion measurements on TAR-G28U reveal back exchange with a GS-like structure**. a.** RD profiles of TAR-G28U U38-N3 and U42-N3 showing the dependence of $R_2 + R_{ex}$ on spinlock power ($\omega 2\pi^{-1}$) and offset ($\Omega_{eff}2\pi^{-1}$). Error bars represent experimental uncertainty (one standard deviation). **b.** Secondary structures of TAR GS and TAR ES2 conformations with populations and exchange rate constant ($k_{ex}$) values from fitting the RD data. **c.** Chemical shift fingerprinting reveals that, based on the two probes for which we have RD data, the TAR-G28U mutant is exchanging with a GS-like conformation.

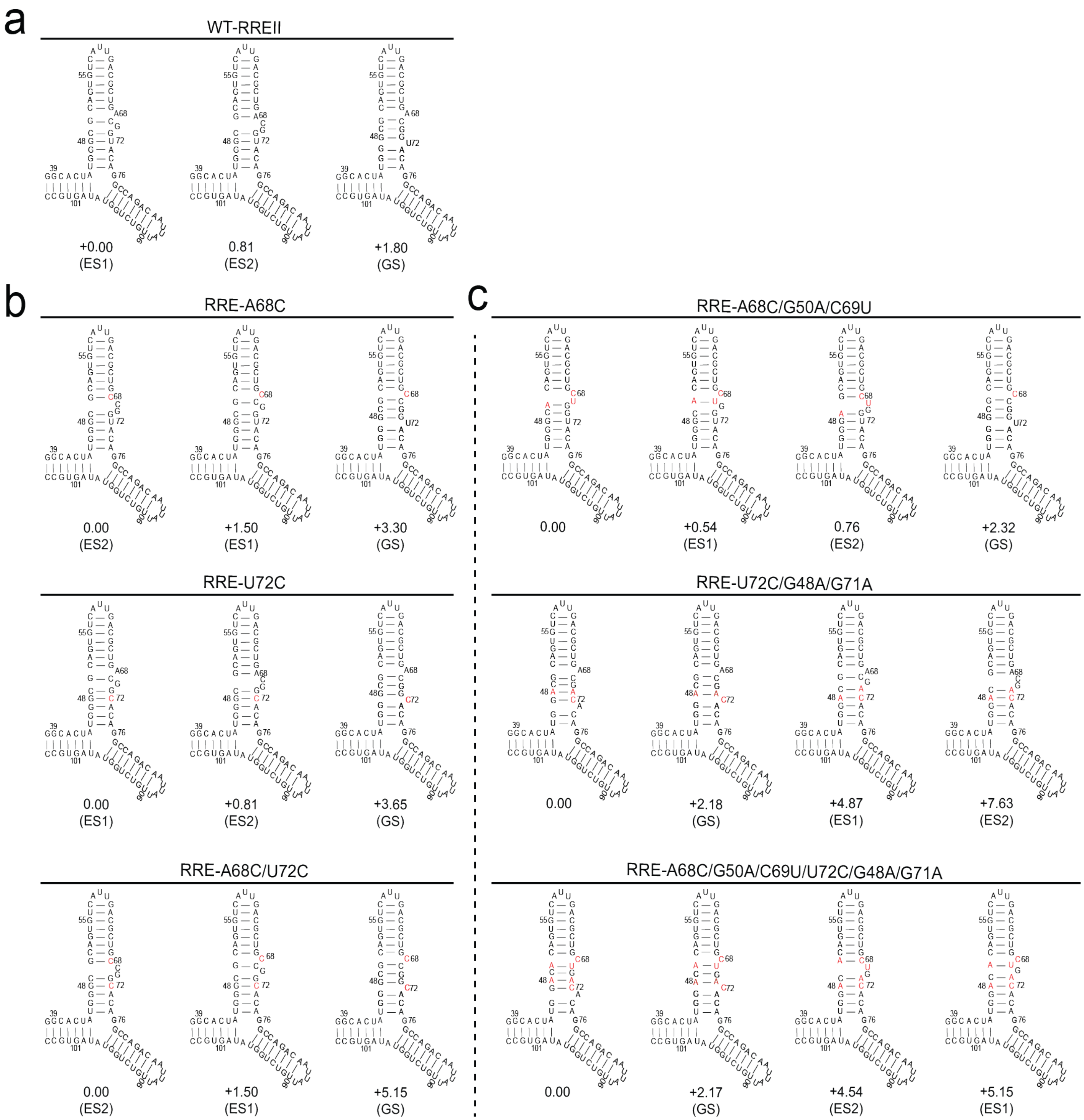


**Supplementary Figure 5:** The energy ranking of trap-and-rescue mutants predicted by MC-fold. The MC-Fold prediction of wtRREII (a) ES-stabilizing (b) and rescue (c) mutations.

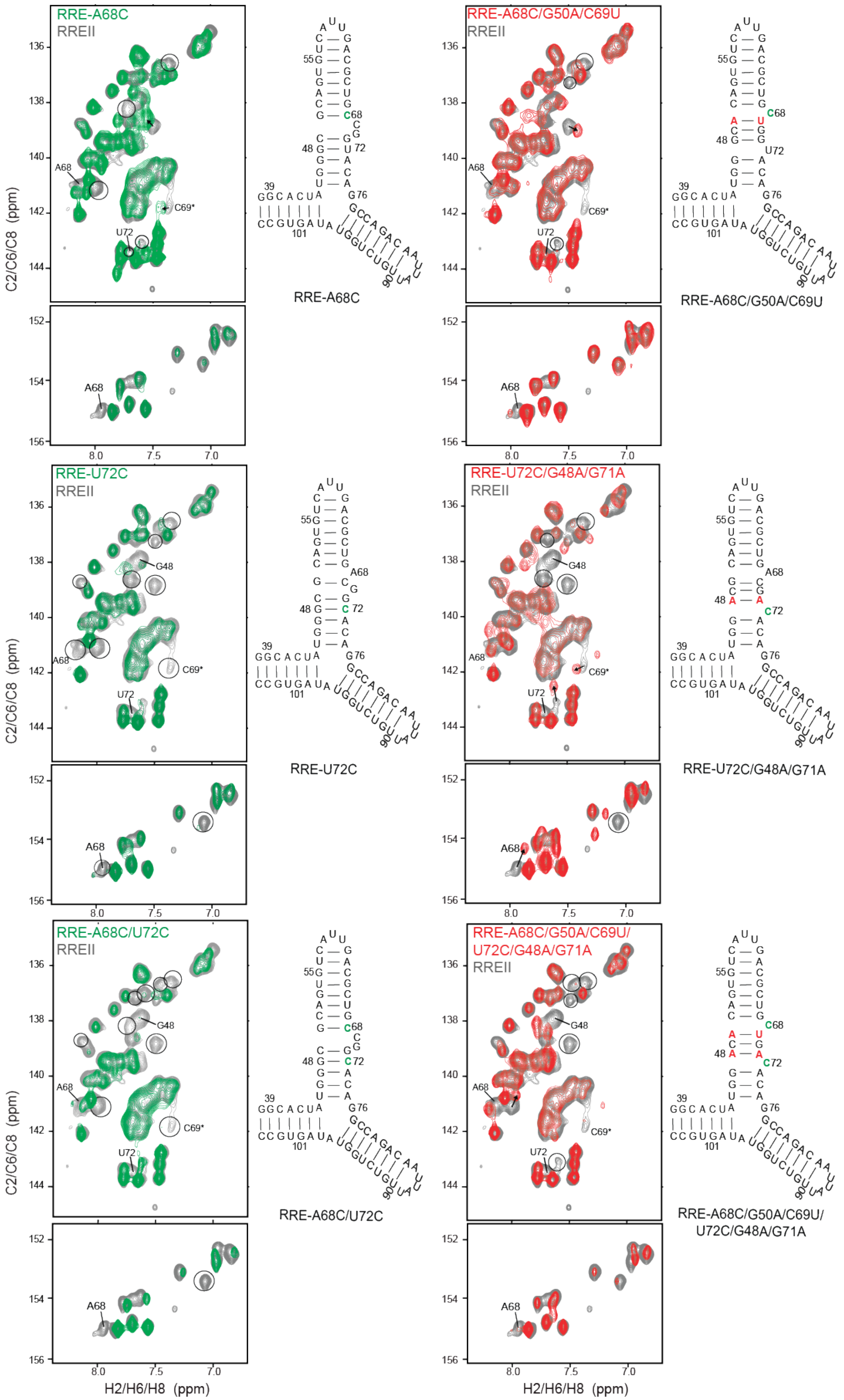



**Supplementary Figure 6:** Chemical shift fingerprinting to test the dominant conformation and extent of stabilization of RRE ES-stabilizing and GS-rescue mutants. RREII ES-stabilizing mutants (green) and rescue mutants (red) overlaid with wtRREII (grey). Resonances that differ with wtRREII are circled, excluding mutated bases.

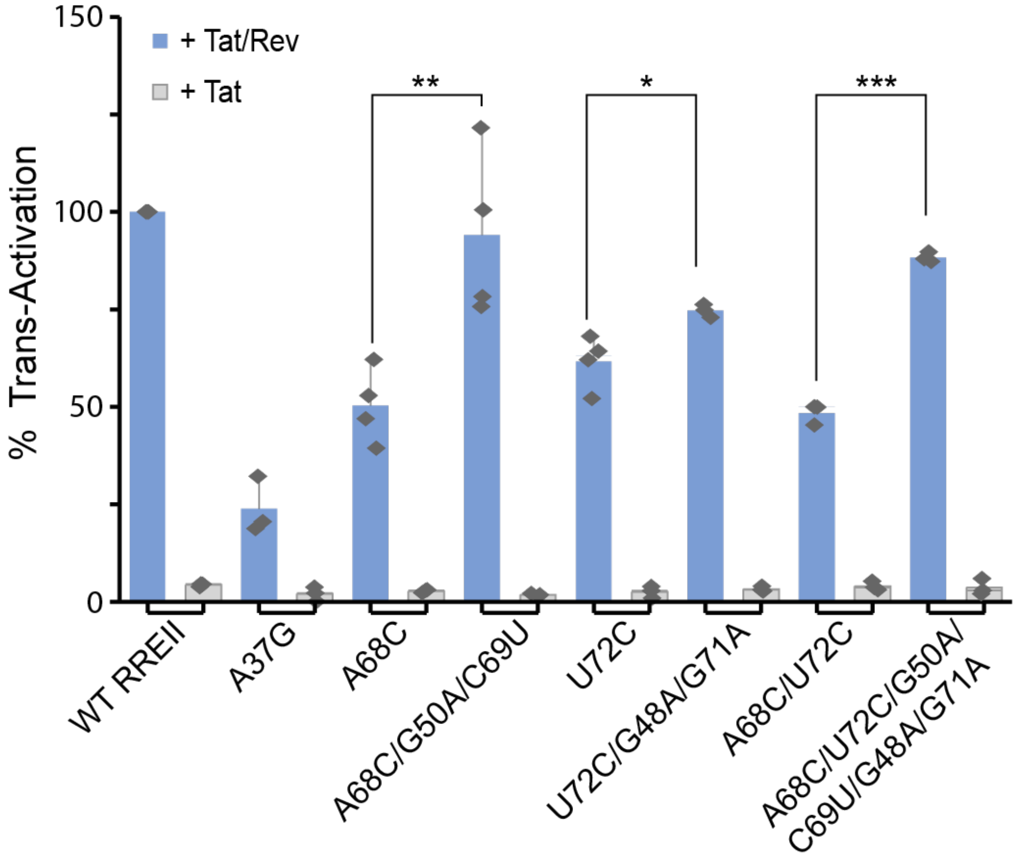


**Supplementary Figure 7:** Cell-based trans-activation assays to detect Rev-RRE binding. Tat-Rev fusion protein dependent trans-activation assays of RRE and mutants. HeLa cells were transfected with pFLuc-TAR/RRE reporter plasmid and an RLuc internal control in the presence of (+ Tat/Rev) or (+Tat) expression plasmid. At 48 hr post transfection, cell lysates were tested for luciferase activity. Reported values are the quotient of FLuc activity and RLuc activity with values normalized to wtRREII for every replicate.

**Supplementary Table 1. Spinlock powers and offsets for R1ρ measurements in TAR-G28U**

| | Off-resonance spinlock power [Offsets] |
|---|---|
| U38-N3 | 100 & ± [10, 32, 64, 96,128,160, 192, 224, 256, 288, 320, 352] |
| | 200 & ± [10, 64,128, 192, 256, 320, 384, 448, 512, 576, 640, 704] |
| | 300 & ± [10, 95,190, 285, 380, 475, 570, 665, 760, 855, 950, 1045] |
| | 600 & ± [10, 191, 382, 573, 764, 955, 1146, 1337, 1528, 1719, 1910, 2101] |
| | 1200 & ± [10, 382, 764, 1146, 1528, 1910, 2292, 2674, 3056, 3438, 3820, 4202] |
| U42-N3 | 100 & ± [10, 32, 64, 96,128,160, 192, 224, 256, 288, 320, 352] |
| | 200 & ± [10, 64,128, 192, 256, 320, 384, 448, 512, 576, 640, 704] |
| | 300 & ± [10, 95,190, 285, 380, 475, 570, 665, 760, 855, 950, 1045] |
| | 600 & ± [10, 191, 382, 573, 764, 955, 1146, 1337, 1528, 1719, 1910, 2101] |
| | 1200 & ± [10, 382, 764, 1146, 1528, 1910, 2292, 2674, 3056, 3438, 3820, 4202] |

**Supplementary Table 2. Exchange parameters obtained from fitting R1ρ data measured for TAR-G28U U38N3**

| | |
|---|---|
| Population (%) | 0.183 ± 0.01 |
| $\Delta\omega_{ES}$ (ppm) | 7.0 ± 0.3 |
| $k_{ex}$ ($s^{-1}$) | 2108 ± 228 |
| $k_1$ ($s^{-1}$) | 3.85 ± 0.46 |
| $k_{-1}$ ($s^{-1}$) | 2103.84 ± 227.49 |
| R1 (Hz) | 1.58 ± 0.03 |
| R2 (Hz) | 8.16 ± 0.12 |
| Reduced $\chi^2$ | 0.42 |
